# Towards the reliable use of aerial eDNA for ecosystem monitoring

**DOI:** 10.64898/2026.05.19.726284

**Authors:** Nadia Sokal, José Ramón Úrbez-Torres, Letitia Da Ros

**Affiliations:** Summerland Research and Development Centre, Agriculture and Agri-Food Canada, Summerland, BC, Canada

**Keywords:** airborne eDNA, biovigilance, ecosystem monitoring, metabarcoding, wide-area monitoring, applied genomics

## Abstract

Evidence supporting the use of airborne eDNA for biodiversity studies and ecosystem monitoring is growing. The promise of wide-area population dynamics data for downstream applications in targeted monitoring of pests and pathogens for agriculture and rare species for conservation is appealing; however, several technical challenges persist. Here, we focused on the development of a comprehensive dataset to facilitate assay development and accelerate the use of aerial sampling for species detection. Year-round metabarcoding data was generated using bacterial, fungal, plant, and arthropod primer sets and resulted in relative abundance estimates for 4,960 amplicon sequence variants (ASVs), 1,748 ASVs of which were assigned to a minimum taxonomic level of genus (bacteria, fungi, plants) or class (arthropods). Sequence diversity assessments and seasonal clustering based on presence/absence detection patterns were performed for individual ASVs, while discerning quantitative changes in seasonal abundance required grouping ASVs to at least the genus level. Examination of the technical aspects of metabarcoding suggested that the use of subsampling allows for consistent detection of genera with relative abundance values above 2 %, even when samples have varying sequencing depths. Sequencing depth was the primary determinate for detecting sporadic and/or rare ASVs. Sampler comparisons, common sources of variation, and the benefits of barcoding regional species to supplement the existing taxonomic databases were discussed. Insufficient knowledge of sampler coverage area for the different organism types was identified as a limitation to the deployment of aerial monitoring networks. Considerations for further aerial metabarcoding efforts are suggested based on our experimental findings.

**Importance:** Our study deals directly with the generation, analysis and limitations of airborne eDNA metabarcoding data for re-use by the broader environmental research community. This includes timing of seasonal detection for possible genera of interest across multiple kingdoms, including bacteria, fungi, plants and animals (specifically arthropods), and support for the generation of local databases to assess the current limitations of universal primers for species/genus taxonomic resolution. With regards to methodology, it continues to build upon established best practices for airborne eDNA collection in areas such as sub-sampling and sampling replicates, sampler type and sequencing depth. To accelerate possible uptake and application of the data, we provide the identified ASVs and their seasonal relative abundances as a resource.

## Introduction

Advancements in sequencing technology continue to increase the resolution at which we can explore the natural world. Enhanced detection of sparse genetic signatures from diffuse media can provide insight into ecosystem species composition, population dynamics, and biodiversity across a wide range of environments (Tournayre *et al*. 2025, Bohmann & Lynggaard 2023, Sullivan *et al*. 2023). These sequence-based surveys have complemented observational analyses, enabling taxonomic classification based on sequence similarity while facilitating the study of rare taxa (Clare *et al*. 2022). This was made possible by the identification of conserved genomic regions that are highly variable between species, such as the 16S ribosomal RNA gene (16S rRNA) for bacteria, the Internal Transcribed Spacer (ITS) sequences in fungi and plants, and Cytochrome Oxidase I (COI) in animals, and the validation of associated universal primers specifically for the purpose of metabarcoding (Klindworth *et al*., 2102, Elbrecht & Leese 2017, Moorhouse-Gann *et al*. 2018, Taberlet, 2012). This type of exploratory work using environmental DNA (eDNA), began with soil microbial communities (Ogram *et al*., 1987, Bowers *et al*. 2011), expanded to aquatic environments (Ficetola *et al*. 2008, Cáliz *et al*. 2018), and more recently has extended to the investigation of airborne eDNA.

Airborne eDNA is a theoretically elegant method for ecosystem monitoring, as it is a non-destructive and non-invasive sampling method. Bioaerosols are collected onto filters using either passive (Atkinson & Roy 2022; Foster *et al*. 2023) or active systems (de Weger *et al*. 2020; Giolai *et al*. 2024) and represent a diverse range of material including suspended microbes, propagules and shed fragments from animals, fungi, plants and microorganisms (Després *et al*. 2012, Tulloch *et al*. 2025). Functional applications of airborne eDNA include biodiversity assessments (Tournayre *et al*. 2025, Bohmann & Lynggaard 2023, Sullivan *et al*. 2023), pathogen detection (Giolai *et al*. 2024), invasive species monitoring (Johnson *et al*. 2021a, Sanders *et al*. 2023), ecosystem changes following restoration efforts (Johnson *et al*. 2021b), and in forensic ecology for determining sample point(s) of origin (Foster *et al*. 2023). The use of airborne eDNA has also been successfully deployed in diverse and complex environments including dense urban areas (Qin *et al*. 2020; Leung *et al*. 2021), temperate and tropical forests (Sanders *et al*. 2023; Garrett *et al*. 2023), and in mixed agricultural regions (Pepenelli *et al*. 2025). Although aerosol sampling has become increasingly common, few studies report data across multiple taxonomic kingdoms within a single study and several challenges related to sample collection, choice of sequencing methods, and data analysis have yet to be optimized (Hoopen *et al*. 2017; Tulloch *et al*. 2025).

Variation in airborne eDNA studies begins with the type of samplers deployed. Use of the different systems is largely dependent on the desired simplicity, cost, required maintenance, flexibility, and longevity of the system needed to address the monitoring objectives (Tulloch *et al*. 2025); although, some controlled comparisons have favored the use of passive airborne eDNA sampling systems (Jager *et al*. 2025). Clarity around the effects of sampling design such as sampling duration, density and what may act as reliable sampling replicates still requires validation (Tulloch *et al*. 2025). The influence of species’ ecology and the different forms of shed DNA on detection have yet to be resolved. From a data analytics perspective, sample processing, sequencing methodology, and the breadth of sequences available in public repositories all influence data resolution. The importance of accounting for contamination in the workflow (Bohmann & Lynggaard 2023), inclusion of a mock community (Banchi *et al*. 2020) and methods to minimizing polymerase biases (Nichols *et al*. 2017) have been previously documented. In airborne eDNA studies, one of two different sequencing strategies is commonly used. Metabarcoding, that targets a specific region of a group of organisms with universal primers, and metagenomics that sequence non-specific regions of all genomes present. For both methods, efficacy should be expected to differ based on the type of organism(s) being detected and the resulting sequencing depth. Adding to the complexity, standardized definitions and terms are actively under review by the broader scientific eDNA community (Tulloch *et al*. 2025), and sequencing terms have been used interchangeably in the past (Taberlet 2012). Lastly, obtaining a positive identification at functional taxonomic levels (family, genus and/or species) depends upon sequence availability within existing repositories and having enough representative sequences to determine the taxonomic resolution a primer set is able to provide. As these repositories are actively adding sequences, it should be expected that many groups of organisms remain underrepresented. Thus far, there have been tremendous efforts by the scientific community worldwide to develop online sequence repositories for biodiversity and population dynamics applications (https://boldsystems.org/; https://www.darwintreeoflife.org/; https://www.ednaexplorer.ca/).

In our metabarcoding study, we examined the use of airborne eDNA for year-round seasonal detection of amplicon sequence variants (ASVs) across multiple taxonomic kingdoms including bacteria, fungi, plants, and arthropods from the same samples. ASV sequence diversity among kingdoms and the taxonomic levels at which changes in seasonal abundance were discernible was assessed. The impact of several technical aspects on the metabarcoding experiment, such as sample replicates, sampler type, sequence depth, and use of a regional sequence database were explored to help guide future metabarcoding efforts. Therefore, we aimed to provide; 1) an investigation of across kingdom eDNA metabarcoding detection capabilities, 2) a comprehensive ASV dataset for use by the community to confirm if and when their organism of interest is detectable using passive airborne eDNA sampling, particularly for use in agroecosystem monitoring, 3) assess if tandem use of regional databases improves genus level detection across kingdoms, and 4) a list of considerations to help address known areas of variability for future airborne eDNA metabarcoding studies.

## Materials and Methods

### Site selection and field setup

Five field sites at the Summerland Research and Development Centre were chosen for eDNA monitoring with passive Spornado (Spornado, Toronto, ON, CA; https://spornadosampler.com/) samplers. Initial experimental design aimed to have a multi-vial cyclone Burkard sampler (Burkard Manufacturing Co, Ltd., Rickmansworth, UK; https://burkard.co.uk/) at each site acting as a control, but power and weather driven maintenance challenges did not allow for reliable sampling from the active Burkard samplers year-round. Spornado samplers were placed along a western field edge and were in proximity to three perennial crops in production, apple, sweet cherry and grape. The sites varied in elevation, natural environment, field maintenance, and pesticide application. Field sites were numbered from highest to lowest elevation. Field site 1 was at the highest elevation in a more secluded area adjacent to natural forest and vegetation. Field site 5 was at the lowest elevation and adjacent to a freshwater creek and a protected provincial park near Okanagan Lake British Columbia, Canada. Field site 3 was centrally located amidst production fields with continuous orchard maintenance and spray applications. Field site 3, as the most accessible, was chosen for the location of additional summer sampling for comparisons of passive Spornado samplers, an ‘active’ Spornado sampler with solar-powered fans allowing for improved air intake and a Burkard active spore sampler over a six-week period. For sampler setup, beginning 10 m in from the field edge, a 1.8 m sampling pole was hammered into the soil to a depth of 30 cm, levelled and allowed to rotate 360° unhindered. At site 3 where multiple Spornado samplers were present, the first sampler was set in 10 m from the field edge with each subsequent sampler separated by a distance of 2 m. The multi-vial cyclone Burkard sampler was inset ∼1 m in from the same field edge (9 m away from the first passive Spornado sampler), leveled and supplied from a continual power source. Sampling from the Burkard occurred at an approximate height of 1m from the ground.

### Sample collection

Spornado cassettes were placed in samplers for eight weeks during each season starting in autumn of 2022 until the end of summer 2023. The eight weeks sampled began on the corresponding equinox or solstice. Each week, cassettes were collected using nitrile gloves and replaced with a new sterile cassette. The cassettes were kept at -30°C until DNA extraction. The Burkard spore trap at site 3 was loaded with 8 x 1.5 mL Eppendorf vials with the timer set to rotate vials once a week. The instrument was checked to ensure vials were successfully rotated, thus corresponding to the weekly sampling being done with the Spornado cassettes.

### Mock community

A mock community was constructed based on local organisms as outlined in Table S1. DNA from all organisms was extracted with the Qiagen DNeasy PowerSoil Pro Kit (Qiagen, Hilden, DE) and pooled at 100 ng for each organism. The mock community was PCR amplified, purified, combined into a next generation sequencing libraries in the same way described below for site samples.

### DNA extraction

Under sterile conditions in a laminar flow hood, the nylon filter was removed from the Spornado cassette with sterile forceps. The filter was placed into tubes from the Qiagen DNeasy PowerSoil Pro Kit. For Burkard samples in 1.5 mL tubes, the PowerBead tube contents were poured into the collection tube under sterile conditions. Instructions from the PowerSoil kit were followed with the exception that samples were processed with the GenoGrinder (Cole-Parmer, Vernon Hills, IL,USA), for 2 x 2 min cycles at 1500 rate with 15 s rest between cycles. A final elution volume of 150 ul was used.

### Pre-library PCR amplification

DNA was PCR amplified using half reactions of the Qiagen PCR Multiplex Kit (Qiagen, Hilden, DE) for the most accurate estimates of relative abundance, although there is a higher associated error rate (Nichols *et al*. 2017). Primers used were designed to have a unique forward and reverse site tag using Oligotag (Coissac *et al*. 2012) and included 1-2 nt heterogeneity spacers to increase primer nucleotide diversity for higher quality Illumina sequencing. This allowed multiplexing of the field sites and inclusion of bacteria, plants, fungi and arthropod specific primers (see Table S2. Five different organism specific primer sets were used; bacteria 16S V3-V4 primers (Thijs *et al*. 2017); fungal ITS2 primers ITS3_KYO1_f and ITS4_r (Chen *et al*. 2022); arthropod 16S primers Chair16S forward and reverse; arthropod COI primers BF3_f and BR2_r (Roger *et al*. 2022); and plant UniPlant ITS2 primers forward and reverse (Moorhouse-Gann *et al*. 2018). Annealing temperatures identified in the above-mentioned studies were used for each of the corresponding primer pairs. Optimal cycle number for PCR amplification was determined by qPCR for each primer set in each season, where 35 cycles was the maximum cycle number to avoid amplification bias (Nichols *et al*. 2017). Each sample had eight identical replicate PCR reactions for each organism primer set. Input template was 1 µl of the extracted total DNA with amplification for 30 cycles with bacteria and fungi primer sets, whereas 2 µl of template with 35 cycles of amplification was used with plant and arthropod primer sets. Cycling conditions were 95°C for 15 min for one cycle; 94°C for 30 sec, annealing temperature for 90 sec, 72°C for 60 sec for primer specific number of cycles; and a 72°C 5-minute final extension. All reactions were prepared in a sterile hood. A 2 ml tube with nuclease-free distilled water was left open in the laminar flow during preparation and afterwards used as negative control template to check for possible contamination of the samples during the preparative steps.

### PCR product pooling and purification

All replicate PCR products were pooled and a 20 µl subset of the pooled PCR products were removed for purification. The HighPrep PCR clean up kit (MagBio Genomics, Gaithersburg, MD, USA) was used for DNA purification and samples were quantified using the Qubit 1x dsDNA HS Assay kit (Invitrogen, Carlsbad, CA, USA).

### Sample normalization

To ensure equivalent sample input into library preparation kits, the samples were normalized according to Qubit values. Samples were pooled by field site, for each primer set target (bacteria, fungi, arthropod 16S, arthropod COI, and plant). Samples were pooled at 60 ng or if the Qubit value was ≤ 2 ng/µl, 30 µl was used. Sites pools were further combined by date. This resulted in 32 final date pools for library preparation that were concentrated to a final volume of 50 µl for library preparation. A final quantification was done to confirm library input.

### Library construction and sequencing

Libraries were constructed using the NEBNext Ultra II DNA PCR-free library prep kit for Illumina (New England Biolabs, Ipswich, MA, USA). Unique Molecular Identifier (UMI) adapters were assigned to each date pool for downstream demultiplexing. The SPRI selection step for adaptor removal was adjusted to use 0.9 x beads to ensure capture of all fragments at 200 bp or larger. The final 32 libraries were quantified using Qubit and pooled in equimolar amounts before final dilution and denaturation. The input library was spiked with 1% PhiX control. The MiSeq Reagent Kit V3 600 cycles paired end kit was used on a MiSeq Sequencing System (Illumina, San Diego, CA, USA).

### Rare species detection and sampler comparisons

To examine rare species detection, Spornado and Burkard samplers from Site 3, from the six summer sampling dates beginning from the solstice (29-Jun-23, 7-Jul-23, 13-Jul-23, 20-Jul-23, 27-Jul-23, 3-Aug-23) were selected for further analysis. Sixteen additional PCR amplification reactions were performed as described above, for each primer set. Three different pools were made from the 16 reactions, composed of three, five and eight PCR replicates. Pooling, normalization and library preparation were performed as described above. For sampler comparisons, two Spornado sampling systems, a passive sampler and an identical sampler with a solar-powered fan were placed 2 m apart at Site 3. Six dates (29-Jun-23, 7-Jul-23, 13-Jul-23, 20-Jul-23, 27-Jul-23, 3-Aug-23) were collected. DNA extraction, pre-library PCR amplification, purification, library preparation, pooling, and normalization were performed as described above. The resulting libraries were sequenced using a MiSeq Reagent Kit V3 600 cycles paired end kit.

### Data analysis and global taxonomic assignment

Data was processed and analyzed using the nf-core/ampliseq pipeline version 2.11.0 (https://zenodo.org/records/14011895), which employs recommended workflows for reliable microbial community data analysis (Straub *et al*. 2020), and executed with Nextflow v24.10.4. Ampliseq is a regularly updated community-curated bioinformatics pipeline (Ewel *et al*. 2020) and for these data, the reproducible software environments from Biocontainers were used (da Veiga Leprevost *et al*. 2017). Each primer set was run individually through the ampliseq pipeline using the same raw fastq file outputs from the MiSeq. Briefly, data quality was assessed with FastQC v0.12.1 (Andrews 2010), summarized with MultiQC (Ewels *et al*. 2016) before being passed through Cutadapt v4.6 (Marcel *et al*. 2011) to trim primers. Trimmed primer sequences were discarded, and sequences that did not contain a primer sequence were considered artifacts. Sequences that passed filtering were then examined as one pool (pseudo-pooled) using R v4.3.2 and the DADA2 package v1.30.0 (Callahan *et al*. 2016) to remove PhiX contamination, discard reads with > 4 expected errors, merge read pairs and remove PCR chimeras. ASVs with lengths < 50 bp were discarded. Taxonomic classification was then performed by DADA2 using Silva 138.1 (Quast *et al*. 2012; McLaren & Callahan 2021) for bacteria 16S amplicons, UNITE general FASTA release for fungi version 10.0 (Abarenkov *et al*. 2024a) for fungal ITS2 amplicons, UNITE general FASTA release for all eukaryotes version 10.0 (Abarenkov *et al*. 2024b) for plant ITS2 amplicons and the COIDB – COI1 Taxonomy Database release 221216 (Sundh *et al*. 2023). While arthropod 16S ASVs were processed and taxonomic classification was attempted using databases such as Genome Taxonomy Database (GTDB), only bacterial classifications were returned and thus this primer set was omitted from downstream analyses. For this reason, the arthropod 16S primer set was chosen to be omitted from downstream sampler comparison and rare abundance analyses. ASV sequences, abundance and DADA2 taxonomic assignments were loaded into QIIME2 v2023.7.0 (Bolyen *et al*. 2019) where ASVs were filtered according to primer pair. For bacterial 16S primers, any ASVs with the taxonomic string containing mitochondria, chloroplast, eukaryota was filtered. For fungi ITS2 primers, no filtering was required as all results were assigned to fungi. For the arthropod COI sequences, mitochondria, chloroplast, chromista, chordata, mollusca, and nematoda were excluded as they were unintended primer targets and the effectiveness of these primers for these taxonomic groupings is unknown. Plant ITS2 primer sequences had mitochondria, chloroplast, fungi, and metazoa assigned ASVs removed. The remaining ASVs were analyzed within QIIME2 where the community data was visualized in a barplot, evaluated with alpha rarefaction curves, had alpha and beta diversity investigated after rarefaction to 1500, 1500, 1250 and 500 counts for bacteria, fungi, arthropod and plant primer sets respectively then used to find differentially abundance in taxa across seasons or samplers with ANCOM-BC (Lin and Peddada, 2020). Rarefaction for comparison of the six weeks of summer sampling were able to be set higher to 1500, 10000, 7300 and 4500 for bacteria, fungi, arthropod and plant primer sets respectively. Rarefaction depths were set according to the organism specific read counts where 90% of libraries were retained.

### Regional plant and mock community sequencing

Tissues were collected in 2 ml tubes from field sites and frozen for storage. Three pre-cooled 3 mm tungsten carbide beads and sand were added and samples ground using the GenoGrinder for 2 x 30 sec cycles at 1500 rpm. DNA was extracted using a modified version of the DNeasy® Plant Mini Kit (Qiagen, Hilden, DE). The ground tissue was first combined with 2% CTAB buffer with RNaseA and incubated for 65°C for 20 min. The DNeasy Plant Mini Kit protocol was used starting at the addition of P3 buffer. PCR was then done for downstream Sanger sequencing of amplicons using the appropriate primers without site tags or spacers to 40 cycles. Then amplicons were purified and quantified. Sanger sequencing was done using 50 ng of prepared amplicon and the appropriate primers according to the organism.

### ASV taxonomic assignment from regional databases

Collaborators provided 16S sequences from Canadian collections and sequencing of regional bacteria. Collaborator, regional plant and mock community sequences were collated and the year round metabarcoding ASV output from the nf-core/ampliseq pipeline was BLAST against the resulting regional database using the Geneious Prime 2025.2.2 (www.geneiousprime.com) ASVs with E values < 10^-5, the highest percent pairwise identity and a sequence length greater than 49 bp were used to identify the best sequence match from the regional database. The taxonomy assigned to ASVs from the regional database was compared to the taxonomy assigned by the Ampliseq pipeline using the global databases at the genus level.

## Results

### Seasonal detection patterns and abundance

All five year-round passive samplers were located within 3 km of each other at the Summerland Research and Development Centre (SuRDC). The surrounding land cover within 10 km of the SuRDC consisted of 34.7 % temperate needleleaf forest, 19.0 % water, 17.6 % temperate shrubland, 17.2 % urban, 7.5 % temperate grassland and 2.6 % cropland (Figure 1A). The British Columbia biogeoclimatic ecosystem classification for the area is PPxh1, indicating a region with Ponderosa pine as the dominant species of mature vegetation, with very dry and hot climatic conditions relative to the rest of the province (Ryan, Lloyd and Iverson, 2022). As samplers were placed within a relatively small monitoring area, overlap of detected ASVs was first examined to assess if each site acted as a sampling replicate. Across all year-round sampling dates, only 19 of the 1520 unique ASVs with assigned taxonomies were common to all five sites (Figure 1). 76% of the 1520 ASVs were unique to a particular sampling location. These sites were therefore treated as subsamples and combined to be representative of the area for improved assessment of seasonal trends. All ASVs and their relative abundances within each weekly sample have been provided in Data S1. ASV detection and taxonomic assignment of the pooled data resulted in 534, 594 and 222 ASVs assigned to genus from bacterial 16S, fungal ITS2 and plant ITS2 primers respectively and 398 ASVs assigned to class using arthropod COI primers for a new total of 1,748 ASVs with sufficient read counts and taxonomic assignments. Four separate phylogenetic trees were generated, one for each kingdom, and visualized to demonstrate the sequence-based diversity present among the resulting ASVs (Figure 2A). Branch length measures of sequence variation were removed in Figure 2B to more clearly depict the branch points and overall groupings of the ASVs in the four trees. Hierarchical clustering based on presence/absence detection of the ASVs throughout the year isolated 11 clusters with notable differences in seasonal patterning (Figure 3A). Clusters with strong detection patterns in summer (cluster 4) or summer and autumn (cluster 1) generally had more equal representation across the categories of organisms. Cluster 2 with consistent detection year-round was enriched with fungal sequences, while clusters 3 (autumn detection) and 8 (spring-autumn detection) consisted predominantly of a mix of fungal and arthropod sequences. Clusters 10 and 11 had consistent detection in the winter season and were enriched with bacteria assigned sequences (Figure 3A). Twenty of the most abundant taxa at each level of taxonomic classification were visualized in Figure 3B. Additional genus level information was provided with the number of assigned unique ASVs indicated at the end of the coloured ribbon, followed by the genus name and the cluster numbers to which the ASVs belong (Figure 3B). Overall trends showed that seasonal patterns of fungal ASVs were variable even within genus, suggesting unique patterns depending on the sequence variant examined, while bacterial detection was more consistent but seasonally dependent (Figure 3A and 3B). For example, although *Antarctolichenia*, *Alternaria* and *Bryoria* were detected year-round, different assigned ASVs had markedly different seasonal patterns and were assigned to different seasonal clusters (Figure 3B; Data S2). *Melampsora* and *Puccina* were enriched in summer and autumn likely matching the various plant hosts. Yeasts within the genus *Phaeococcomyces* were not present in the winter, while *Vishniacozyma* were enriched in the winter and spring. Bacterial genera such as *Pedobacter* and *Pseudomonas* were broadly present year around while *Dyadobacter*, *Flavobacterium* and *Chryseobacterium* were present almost exclusively in the winter months. *Nocardioides* were enriched in spring, summer autumn seasons while *Segetibacter* were enriched in summer and autumn. For the abundant plant representatives, *Pinus* were detected in spring/summer, *Amaranthus* in summer/autumn and *Artemisia* detection more prevalent in the autumn season. Log fold change in abundance for taxa with sufficient abundance and detection data was performed using ANCOM-BC with results presented in Figure 4. This confirms trends observed in the presence/absence data, but highlights the taxa with statistically significant seasonal patterns as opposed to those with the highest number of sequencing reads.

**Figure 1:**
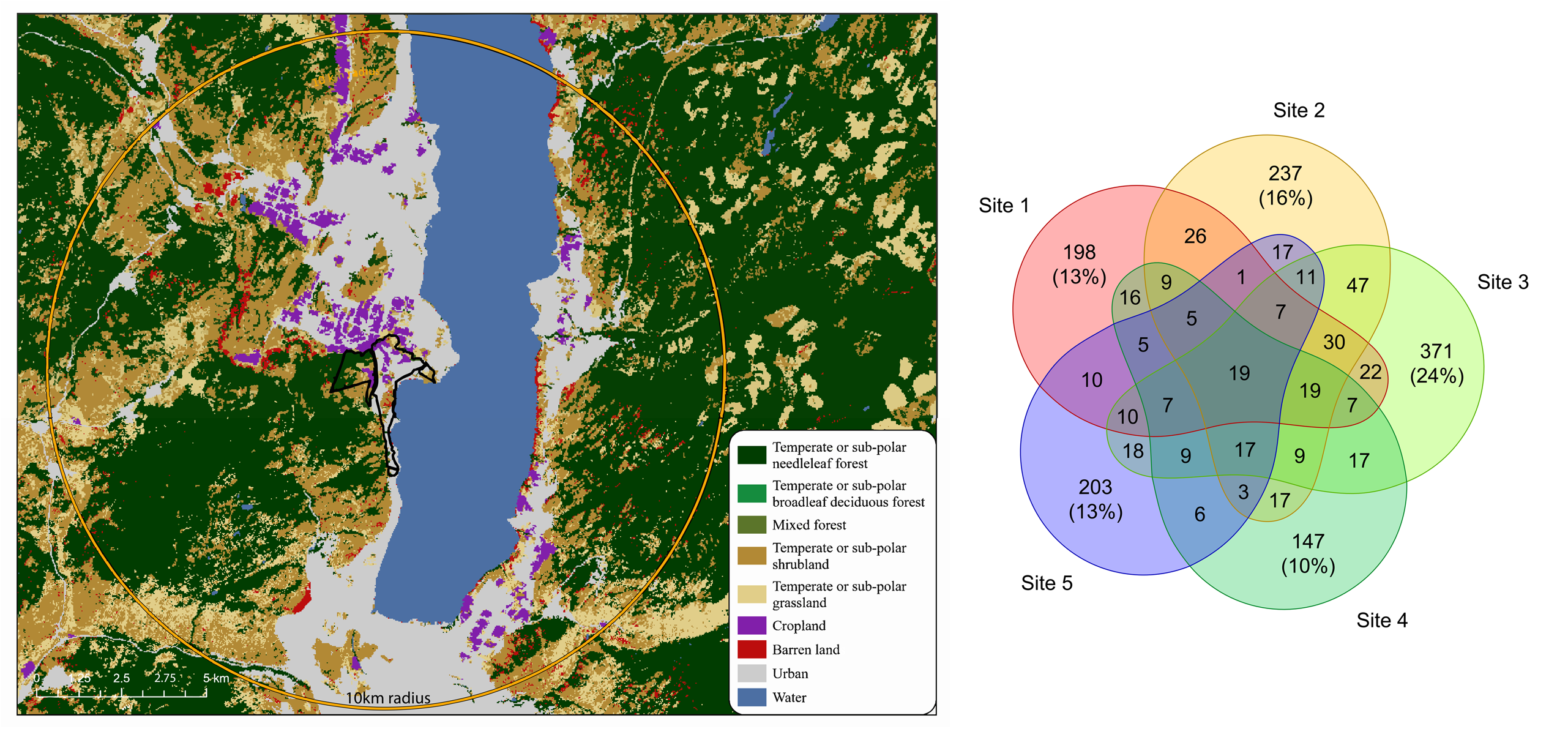
A) Land cover map of the area surrounding the Summerland Research and Development Centre indicated by the black outline. All five sampling sites were located within the boundaries of SuRDC. Land cover composition within a 10 km radius was 34.71% needleleaf forest, 19.02% water, 17.58% shrubland, 17.22% urban, 7.48% grassland, 2.60% cropland and 1.39% barren land. B) Venn diagram showing the overlap in detected ASVs from year-round passive air sampling at five sites in Summerland. A total of 1520 ASVs were used in the comparison. ASVs included those that were identified to genus for bacteria (475), fungi (573) and plants (178) and to class for arthropods (294). All sites were within 3 km of each other next to horticulture production fields. Primary differences between the sites were elevation and land cover adjacent to the production areas. Sites were numbered based on elevation with 1 being at the highest elevation and 5 at the lowest.

**Figure 2:**
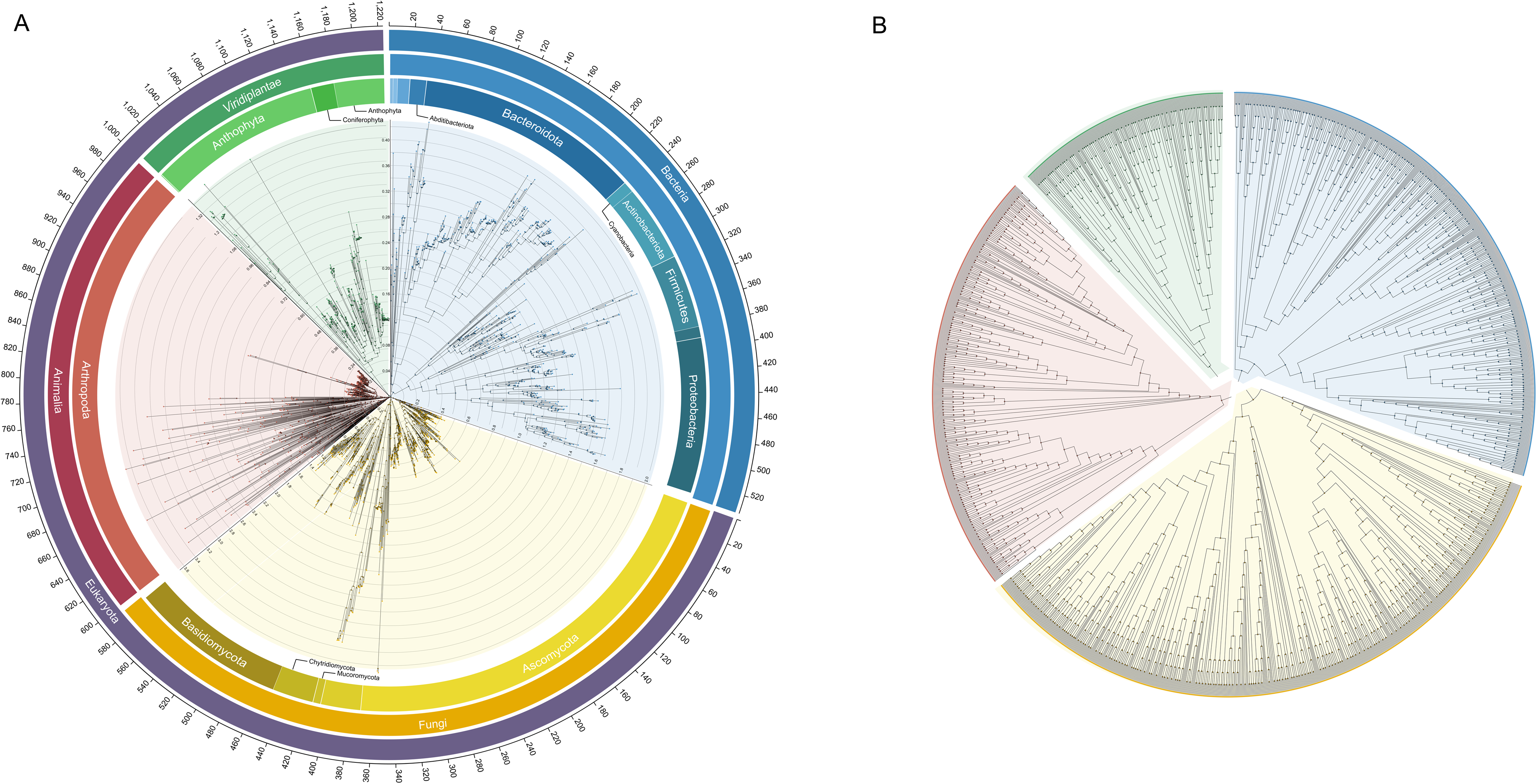
Taxonomic classification of the 1,748 amplicon sequence variants (ASVs) that were identified during year-round passive air sampling to the level of genus for bacterial 16S, fungal ITS2, and plant ITS2 primer sets organized as four separate phylogenetic trees. For the arthropod COI primer set, those identified to class are shown above as only 15 of the 398 were able to be assigned to a lower taxonomic level. ASV counts from each primer set were 534, 594, 222 and 398 for bacterial, fungal, plant and arthropod sets respectively. A) Outer scale on the circle denotes the counts of ASVs with the subsequent rings indicating the domain, kingdom and phylum of the ASVs in the tree leaves below. Phylum is approximate as the inner phylogenetic trees are sequence-based. Each kingdom has an individual phylogenetic tree with a different scale to best visualize the branch length variation. The scale comes before the corresponding tree when traveling clockwise around the graph. Branch lengths in the phylogenetic trees reflect sequence divergence where numeric values indicate average number of substitutions per site. Leaf nodes are highlighted with a marker colour according to kingdom and internal nodes where branching points occur are denoted by a black outlined marker. B) The same phylogenetic trees depicted in A but with branch lengths ignored to more clearly show the branching structure within the trees. Colour of the leaf node indicates the kingdom to which the ASV was classified.

**Figure 3:**
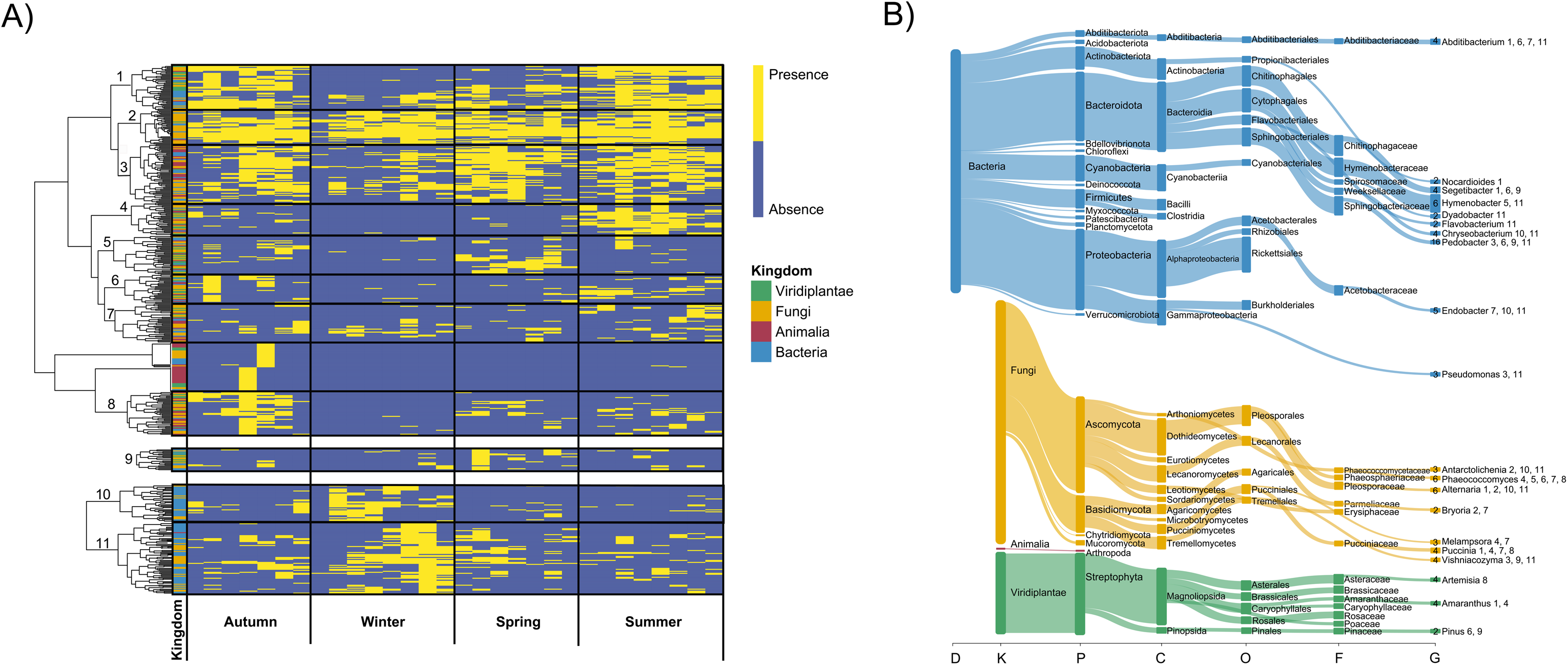
A) Presence and absence heatmap across the full year of passive spore sampling for the 1,748 ASVs with sufficient taxonomic classification. Clusters with definitive seasonal patterning were assigned numbers, isolated and depicted above. A total of 460 ASVs were in the eleven numbered clusters. The full heatmap is provided as Figure S1. ASVs were hierarchically clustered using binary distance with Ward’s D2 linkage method. B) The twenty most abundant taxa at the various taxonomic levels as determined by kraken2 and visualized using Pavian. Numbers to the left of the genus name indicate the count of ASVs that were in one of the eleven clusters. Numbers to the right of the genus indicate to which clusters the ASVs belonged. Data associated with the 460 ASVs has been provided as Data S2.

**Figure 4:**
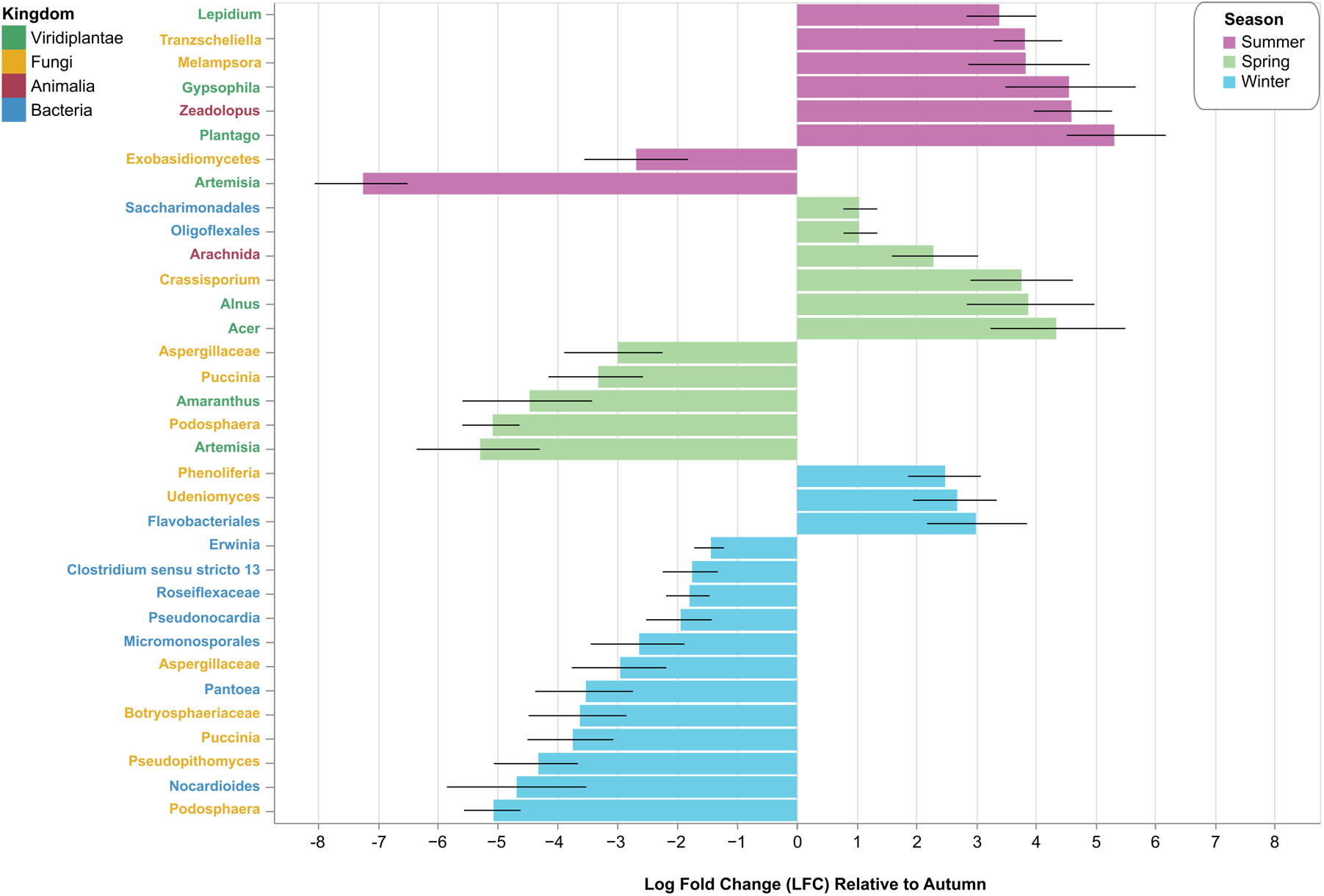
Differential seasonal abundance for taxa on which analysis of composition of microbiomes with bias correction (ANCOM-BC) could be applied. Results are displayed as log fold change relative to the autumn season and only include those with statistically significant changes across seasons. Names of the various taxa are coloured according to the kingdom to which they belong.

### General technical considerations

Airbourne eDNA analysis is in the early stages of implementation for use as a monitoring tool, particularly for short-term monitoring where cost-effectiveness, ease of use, maintenance and reproducibility are major considerations, each with broad impacts on downstream use. To help inform future implementation of airborne eDNA, we outline below a series of smaller experimental outcomes. In the initial seasonal field sampling setup, Burkard spore traps were intended as the industry standard for active air sampling alongside the commercially available Spornado passive spore samplers. However, power requirements and weather-induced mechanical failures in autumn and winter led to inconsistent sampling and sample contamination with the more mechanically complex Burkard samplers. Additionally, in spring and summer months, smaller invertebrates were pulled into the active sampler itself. These are sources of detection error and should be carefully considered when choosing samplers for deployment for a particular study or application. A second source of possible detection error would be DNA contamination during the workflow. For this study, all samples were processed for the DNA extraction and PCR amplification reactions in a laminar flow hood alongside an aliquot of water that was left open throughout the processing. The water was PCR amplified and sequenced. Few fungal and bacterial ASVs were identified from the open water samples, and no evidence of contamination was observed in any of the seasonal, sampler or rare abundance libraries. Although some ASVs from the open water control were also detected in the environmental samples, it was not consistently observed in a group of samples processed within the same batch, resulting in its use as a true observed value rather than a contaminant of concern. Verification of processing contaminants is a requirement of airborne eDNA sampling as very low DNA concentrations were initially extracted. Examples from a subset of summer collections showed concentrations ranging from 0.13 to 1.47 ng/uL from passive Spornado samplers and 2.42 – 5.24 ng/uL recovered from Burkard active sampling. Differing amounts of extracted DNA were not found to be predictive of number of ASVs detected. Using the same summer samples, increasing the number of PCR reactions pooled per library preparation was not found to have an effect on the detection of rare ASVs. Pooling 3, 5, or 8 PCR reactions did not reliably or consistently result in higher ASV detection for either Burkard or Spornado samplers. Sequencing depth and subsampling were major factors in the detection of rare ASVs and were explored in more detail.

### Detection variation among samplers

Site 3 was chosen for additional experiments to compare sampler effectiveness across the multiple primer sets (bacteria, fungi, plant, arthropod). Samplers included a Burkard spore sampler, two passive Spornado samplers and one Spornado sampler with a solar powered fan for a total of four samplers. The experiments were initially structured as two paired experiments (Burkard vs. passive Spornado, passive vs solar), but as weekly collections, location and sequencing run were identical, all samplers were compared. Number of ASVs detected varied from 1411 to 3321 with a difference of 1303 ASVs between the two passive samplers located at the same site. 714 ASVs were detected by all four samplers with 1218 shared between the two passive samplers (Figure 5). The passive Spornado sampler paired with the Burkard had significantly greater diversity of bacteria and arthropod assigned ASVs. However, greater examination of the large differences between identical samplers at the same site warranted further investigation to confirm if sampling replicates were possible to achieve when conducting airborne eDNA studies (Figure 5). No benefits were observed from the addition of the solar fan to the passive sampler in this study.

**Figure 5:**
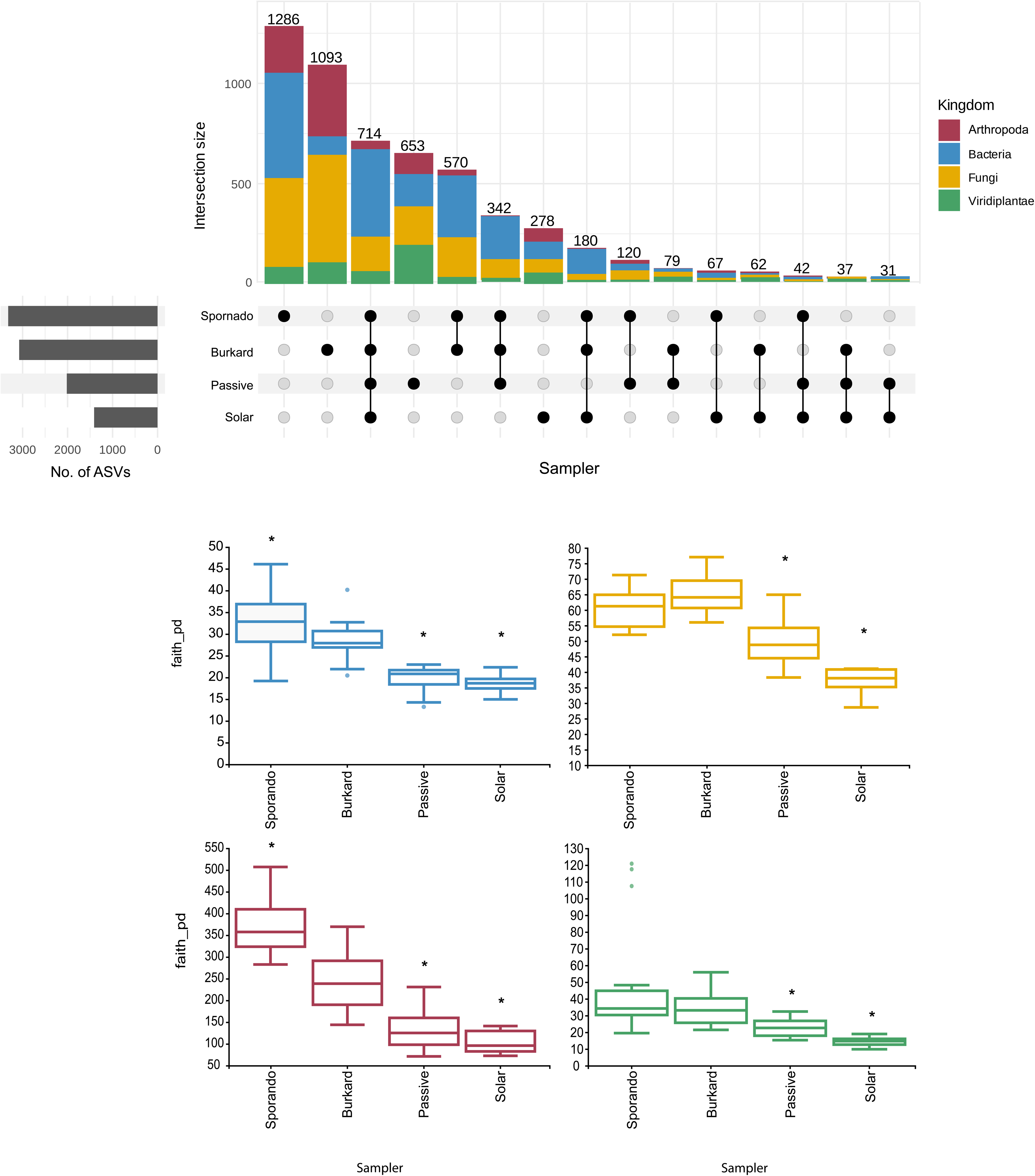
A) Upset plot demonstrating the overlap in detection of ASVs for those classified to genus for bacteria, fungi and plants and class for arthropoda. All samplers were placed at Site 3 from weeks 1 to 6 of the summer season (June 21 to August 3). Set size indicates the total classified ASVs across all groups per sampler. Boxplots are faith’s phylogenetic diversity measurement of alpha diversity for each of the primer pairs to compare performance of the respective samplers. Significant differences compared to the Burkard at a p-value of < 0.05 is denoted by an asterisk above the corresponding boxplot.

Variability in ASVs detected by identical passive samplers at the same site led to additional comparisons to identify ways in which the effectiveness of airborne eDNA sampling methods could be improved. Comparisons were made between a new passive sampler used only for the six weeks of sampling (1S-1) acting as the control, the second Site 3 passive sampler (1S-2) that had been used for sampling throughout the year, and the lower depth sequencing of the pooled 5 sites during the same six weeks of summer (5S-1) in the same year. Results suggest that rarefaction could not entirely eliminate effects of differences in sequencing depth between S1 and S5 samples. However, even samples 1S-1 and 1S-2, which were experimental replicates sequenced in the same run, showed significant differences in calculated alpha- and beta- diversity metrics. Interestingly, although samples 1S-1 and 1S-2 had higher taxonomic richness than the low- depth pooled sample 5S-1, all three groups had similar evenness, suggesting the additional taxa was not substantially altering the overall structure of the existing community. This is reflected in the relative abundance patterns of taxa across samples for each primer pair (Figure 6A; Data S3). Using a range of summed six-week relative abundance thresholds, the percentages of 1S-1 genera detected in 1S-2 and 5S-1 were calculated. All datasets detected the most abundant taxa (> 2 % summed relative abundance), with ≥ 90% of control 1S-1 genera detected by all samplers (Figure 6B). 1S-1 and 1S-2 had detection similarities greater than 75% down to a six-week summed relative abundance threshold of 0.01% across bacterial, fungal and plant primer sets. The fewest similarities were at the lowest abundance threshold for the plant primer set (Figure 6). The arthropod COI primer set was not included in the assessment as there was not sufficient taxonomic assignment of the ASVs beyond Class (Figure 6A). This suggests that airborne sampling at the same site can function as a replicate, but sequence depth should be as consistent as possible for taxa with less than 1 % summed relative abundance.

**Figure 6:**
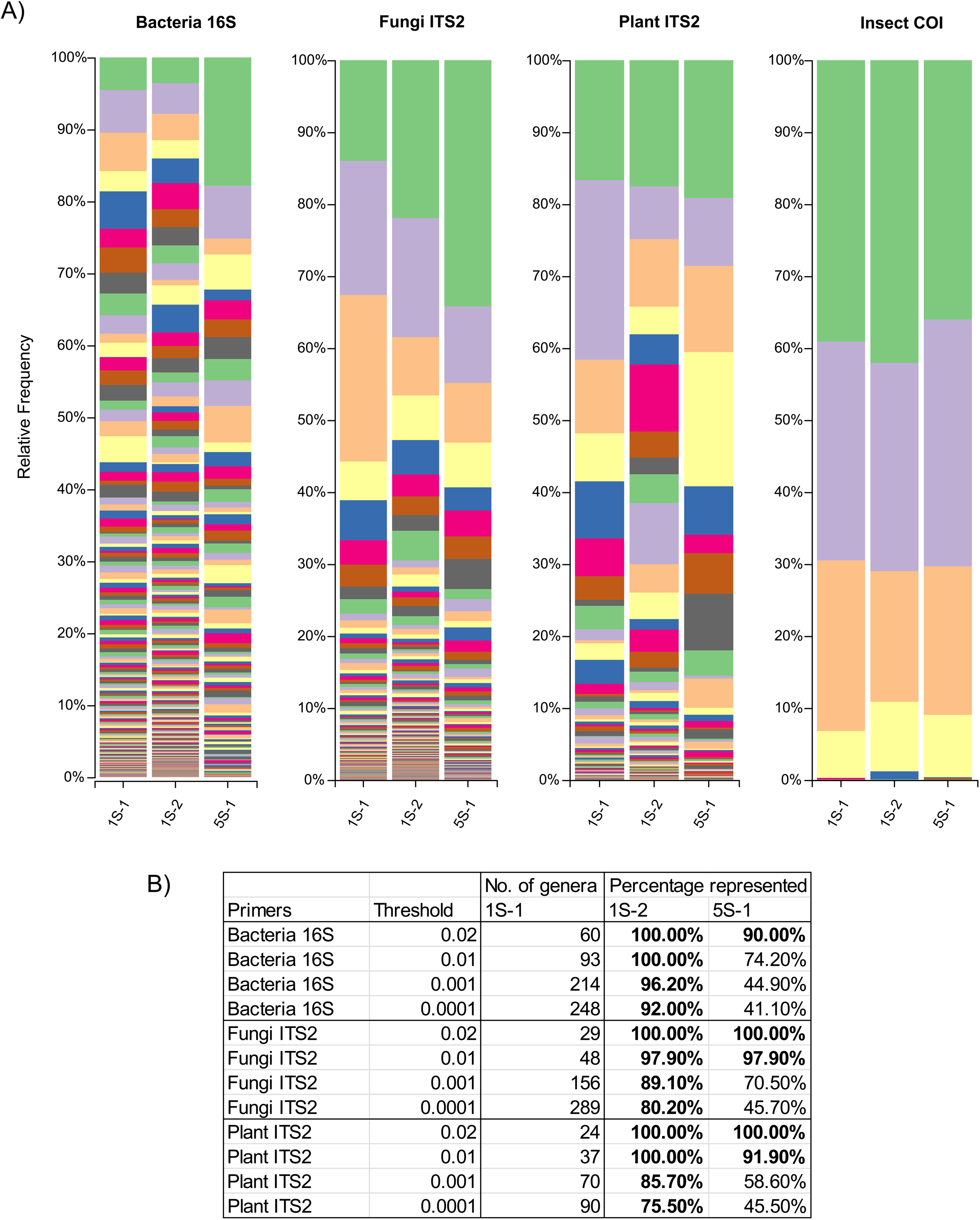
Visualization of diversity within and across passive samplers according to taxonomic assignments. Data from weeks 1 through 6 in the summer season were used. Site 3 - Summer is a passive sampler that was new and used only for the 6 weeks in the summer. Site 3 - year round is a second passive sampler locate ∼2 m from the first that was used for weekly sampling all year round although only summer weeks 1 -6 were PCR amplified again for this comparison. All sites - year round was the pooled summer weeks 1-6 from all five sites but sequenced to a lower depth. In the table, taxa detection was compared at different summed relative abundance thresholds across the 6 weeks using Site 3 - Summer as the control. Taxa detection similarity above 75% are bolded.

### Taxonomic assignment of ASVs from regional databases

Our regional databases were sequences from local collections completed during this study (Data S4; plant ITS2 sequences) or those provided by collaborators (bacteria 16Sv3-v4 sequences) that were collected in Canada. In total there were 7053 and 145 unique regional sequences for use in taxonomic identification in bacteria and plants respectively. Comparisons were made at the genus level to see if regional and global databases returned the same best match using unique ASVs from the year-round data (Data S1) as input. From the combined global and regional sequence databases, 579 of the 736 bacterial ASVs and 247 of the 282 plant ASVs were assigned a putative genus. Of the ASVs that returned a putative genus from both the global and regional datasets, 79.5% and 27.6% were assigned same genus for bacteria and plant ASVs respectively. Overall, the inclusion of the regional bacteria and plant datasets increased the total number of ASVs with genus-level taxonomic assignments by 6.3% and 8.9% respectively. Similar comparisons were attempted with regional fungal and insect datasets, however insufficient numbers of sequences were available for comparison (< 50 sequences per dataset).

## Discussion

The dispersal potential of organisms shapes both species population dynamics and the overall structure of ecological communities. Wind is a key dispersal mechanism with evolutionary adaptations found in all branches of life. Size, shape, timing and weather patterns determine spore dispersal processes, seed and pollen transmission, insect dispersion and response to changing environmental conditions as well as the atmospheric dispersion and deposition of microbial communities (Naranjo-Orrico *et al*. 2025; Tackenberg *et al*. 2003; Reynolds *et al*. 2017; Santl-Temkiv *et al*. 2022). While representing a promising arena for ecosystem-level monitoring of biodiversity, pests, pathogens and disease, airborne eDNA sampling has inherently high variability. Shifting environmental conditions, diffusely distributed DNA and species dispersal characteristics are unique challenges of airborne eDNA studies and have broad impacts on the results (Tulloch *et al*. 2025). Yet it is well documented that aerial metagenomics and metabarcoding is a valuable tool for tracking ecosystem diversity and taxa of interest in terrestrial ecosystems (Roger *et al*. 2022; Lynggaard *et al*. 2022; Sullivan *et al*. 2023; Giolai *et al*. 2024; Tournayre *et al*. 2025). Here we aimed to employ best practices in metabarcoding for accuracy in relative abundance estimates (Nichols *et al*. 2017) and reduced variation from known technical sources of error (Tulloch *et al*. 2025). This was done by using the nf-core/ampliseq pipeline, limiting contamination of DNA, and by assessing the effects of parameters such as sequencing depth, varying levels of DNA capture, and changes in ASV taxonomic assignment resulting from querying both a global and regional sequence database. The major sources of technical variation were sequencing depth and designation of a sampling replicate. Higher depth sequencing significantly increased the number of taxa detected but did not significantly alter community structure (Figure 6). Added taxa were predominantly those of very low abundance, possibly a result of the efforts made to minimize polymerase and amplification effects on abundance results. Overall, the use of methodology that preserves relative abundance of the original sample can allow for reliable seasonal detection of ASVs across varying sequencing depth, however higher sequencing is essential for reliable detection of sporadic or rare ASVs. Variation in taxa detection across sites was higher than expected given site proximity (Figure 1) and subsampling within a region appears to be an effective way of increasing number of detected ASVs. To our knowledge, few examinations of sampler coverage (radius in km) and the effectiveness of interpolate of detection between sites have been done and requires further validated prior to wide-scale adoption of aerial wide-area monitoring (Tournayre *et al*. 2025).

eDNA studies have largely focused on a single organism group, however there is a growing interest in simultaneous detection and analyses of multiple kingdoms (Banchi *et al*. 2020; Sullivan *et al*. 2023). Here we employed four primer sets, bacterial (Thijs *et al*. 2017), fungal (Chen *et al*. 2022), plant (Morrhouse-Gann *et al*. 2018) and arthropod (Roger *et al*. 2022) primers to obtain a well-rounded dataset of interest to many fields of study, so it may be leveraged for downstream ecosystem monitoring applications. Seasonal trends observed in this study echo those found in other airborne studies in temperate regions (Banchi *et al*. 2020; Sullivan *et al*. 2023; Abrego *et al*. 2024). Fungal ASVs had the highest sequence diversity and second highest detection rates throughout the year (Figure 1A). While many fungal genera had abundant reads, differential seasonal abundance was not possible likely to the high variability in individual ASV detection patterns (Figure 3). To highlight this point, fungal genera with measurable differences across seasons were predominantly restricted to fungal plant pathogens or cold-adapted fungi (Figure 4). Bacterial genera had more consistent seasonal patterns, however detection was more concentrated in the winter months (Figure 3). In terms of abundance, 68.1% percent of reads assigned to bacteria were detected during the eight weeks in winter, compared to the 8.7 and 8.9% of reads detected in spring and summer respectively. Of the 146 bacterial ASV assigned to genus, 126 were detected in winter with 22 uniquely found in the winter season. Between 82- 90 of the 146 ASVs were found in spring summer or autumn. While this is in agreement with previous reports indicating air temperature has significant impacts on the airborne bacterial community (Genitsaris *et al*. 2017; Sullivan *et al*. 2023), it does refute some suggestions that highest bacterial abundance can be found during the summer months (Genitsaris et al. 2017). A combination of seasonal wind direction, ultraviolet radiation, humidity and emission intensity within a biogeoclimatic zone likely determine the timing of peak bacterial abundance (Gong *et al*. 2020; Wang *et al*. 2025). Genera that may aid in elucidating key meterological parameters affecting microbial community structure include common soil plant and water bacteria such as *Arthrobacter, Brachybacterium, Massilia*, *Pedobacter*, *Pseudomonas*, *Sedminibacterium*, *Sphingomonas* as they were generally abundant and detected year-round. Human-associated bacteria (e.g. *Blautia*, *Catenibacterium Collinsella*, *Dorea*, *Holdemanella*, *Peptoclostridium* and *Staphylococcus*) were enriched in the winter. Further seasonal studies on bacterial and fungal viability would be an interesting addition to metabarcoding datasets (Gong *et al*. 2020).

Data associated with bacteria and fungi were more amenable to statistical analyses as there were more abundant read counts, more consistent ASV detection and comprehensive metabarcoding databases for these two organism groups (Quast *et al*. 2012; McLaren & Callahan 2021; Abarenkov *et al*. 2024a). Limitations to data analysis with plants and arthropods are greater in that the universal primers are not as well characterized, comparatively speaking aerial dispersal is more dilute and infrequent relative to fungi and bacteria, community structure is more geographically biased and metabarcoding databases are currently under-development (Arstingstall *et al*. 2021; Salis *et al*. 2024). The importance of having confirmed taxonomic assignments for metabarcoding sequences was most apparent when examining the plant ITS2 results. Global and regional databases gave matching taxonomic assignments for only 27.6% of plant ASVs identified to the genus level from the seasonal year-round data and furthermore species level data were not congruent. An example of this is that the *Pinus* genus was assigned to 15 unique ASVs using global database with an additional 2 ASVs assigned after supplementing with the regional database sequences. However, none of the species designations matched between the global and regional datasets. The global dataset returned *Pinus contorta*, *Pinus hartwegii*, *Picea rubens* and *Keteleeria fortunei* for ASVs that all matched to *Pinus ponderosa*, the local dominant pine species, using the regional sequence data. A similar scenario was found with the *Prunus* genus where the five ASVs with regional matches to *Prunus avium* were identified as *Prunus sunhangii* and *Rosa bracteata.* This is further complicated by the knowledge that in the orchards surrounding the sampling sites, many other *Prunus* species are present including; *Prunus cerasus, Prunus virginiana, Prunus persica, Prunus armeniaca,* and *Prunus domestica*. Lack of species level sequence data affects our ability to determine to what taxonomic level these conserved regions can differentiate, is needed to set a threshold sequence fragment length for accurate genus level assignment and inhibits our ability to merge abundance data of ASVs that are likely to be detecting the same species. Particularly for kingdoms with fewer curated barcoding sequences, as is the case for plants and insects. The generation of supplementary regional databases from local collections could incrementally increase resolution for species of interest, particularly if the resulting sequences are then deposited into existing public repositories. To that same point, greater number of sequenced representatives within a genus determines whether the use of universal primers and wide-area airborne eDNA datasets can be used for species level resolution or if subsequent species-level assays from airborne eDNA samples are required as can be the case with fungal phytopathogens (Chen *et al*. 2024). Species-level resolution is needed to unravel which physical and physiological traits allow for aerial detection across seasons. For example, using some of the plant genera with the highest detected abundance *Artemesia*, *Amaranthus* and *Pinus*, points of detection varied by ASV throughout the autumn, spring and summer. Of the twelve ASVs assigned to *Amaranthus*, seven were detected in autumn, eight in summer and two in spring and five with multi-seasonal detections. For the 25 ASVs assigned to *Artemisia*, there are high relative abundance values in the autumn, although the length of time an ASV was detected varies widely. The 15 *Pinus* ASVs had more sporadic detection in late spring and summer. It remains unclear as to why some ASVs within a given taxa show these patterns of prolonged or multi-seasonal detection (Data S1). Without greater resolving of the data, validation of the effects of species biology and dispersal mechanisms on airborne detection would prove difficult. This highlights the importance for careful consideration of what exists locally, what the detection goals are for experiments and the need for submission of sequences to public repositories to enhance the reliability of monitoring applications using airborne eDNA.

Taken together, the findings of this study were used to inform the following recommendations for future aerial metabarcoding efforts: 1) the amount of multiplexing should be limited to improve rare ASV detection over other technical adjustments, 2) passive samplers provide reliable detection and can be advantageous when sampling across seasons or for larger scale deployments, 3) consider subsampling within 3 km for each area of interest for a more representative observation of community structure and variation, 4) consider building a regional database for plant and arthropod species to enable higher resolution taxonomic identification of ASVs and enable possible characterization of species traits that affect the likelihood of detection by aerial sampling. If focusing on particular pathogens of interest, curating a regionally focused dataset for bacteria and fungi would be beneficial to assess expected sequence diversity within the conserved genomic regions and identify ASVs that could be merged for more accurate measures of relative abundance.

## Conclusion

Airborne eDNA has potential for wide area monitoring of population dynamics for informed management of agroecosystems, early detection of invasive species and climate impacts on biodiversity. To accelerate possible uptake and application, we provide the identified ASVs and their seasonal relative abundances as a resource for first assessments as to if a given taxa of interest could be detected as well as what time of year sampling efforts may be focused. The use of aerial sampling in tandem with on-the-ground monitoring efforts would further explore the effectiveness of the approach, with the goal of improving our response to evolving threats to food production and the preservation of natural ecosystems.

## Data availability statement

Year-round and sampler comparison nf-core ampliseq qiime2 data outputs of all ASVs and their relative abundance are provided as supplement files. ASVs whose IDs have been classified to at least a taxonomic level of genus and those verified using local databases aim to be submitted to the eDNA Explorer Canada platform for broader accessibility.

## Acknowledgments

The authors thank Julia Douglas-Freitas for their technical support in implementing the nf-core ampliseq pipeline on the Shared Services Canada General Purpose Science Cluster (GPSC), Dr Jennifer Town for use of their regional 16S sequences, Molina Tek for the land classification map and land cover statistics, Arlan Benn for direction on field sampling considerations, as well as Jordan Fraser, André Rachinski, Talia Morfopoulos and Rebecca Saunders for their help with field sample collections.

## Author contributions

L.D. and J.U.T. conceived the study and obtained funding. J.U.T. managed the project. L.D. designed and implemented the study. N.S. performed the sample preparation and sequencing. L.D. analyzed the data and generated the figures. N.S. analyzed sequences against regional databases. L.D. and N.S. co-wrote the manuscript. L.D. and J.U.T. edited and finalized the manuscript. All authors read and approved the manuscript.

## Supplemental Information

**Figure S1.** Complete presence/absence heatmap from year-round passive spore sampling data with the 1,748 ASVs across the four primer sets with sufficient taxonomic classification (genus for bacteria, fungi, plant and class for arthropods).

**Table S1:** Organisms collected and used for the mock community

**Table S2:** Primers used for PCR amplification

**Data S1.** Relative abundance and taxonomic classifications of ASVs from year-round sampling DADA2/QIIME2 outputs.

**Data S2.** The 460 ASVs found in the eleven designated clusters and relative abundance within each week throughout the sampling seasons.

**Data S3.** Relative abundance and taxonomic classifications of ASVs from sampler comparison DADA2/QIIME2 outputs.

**Data S4.** Sanger sequences generated using ITS2 plant primers for the regional database from local plant species.

